# Genotypic variation in blueberry flower morphology and nectar reward content affects pollinator attraction in a diverse breeding population

**DOI:** 10.1101/2024.02.22.581568

**Authors:** Juliana Cromie, John J. Ternest, Andrew P. Komatz, Paul M. Adunola, Camila Azevedo, Rachel E. Mallinger, Patricio R. Muñoz

## Abstract

Pollination is crucial to obtain optimal blueberry yield and fruit quality. Despite substantial investments in seasonal beekeeping services, blueberry producers consistently report suboptimal pollinator visitation and fruit set in some cultivars. Flower morphology and floral rewards are among the key factors that have shown to contribute to pollinator attraction, however little is known about their relative importance for improving yield in the context of plant breeding. Clarifying the relationships between flower morphology, nectar reward content, pollinator recruitment, and pollination outcomes, as well as their genetic components, can inform breeding priorities for enhancing yield. In the present study, we measured ten flower and nectar traits and indices of successful pollination, including fruit set, seed count, and fruit weight in 38 southern highbush blueberry genotypes. Additionally, we assessed pollinator visitation frequency and foraging behavior over two growing seasons. Several statistical models were tested to optimize the prediction of pollinator visitation and pollination success, including partial least squares, BayesB, ridge-regression, and random forest. Random forest models obtained high predictive abilities for pollinator visitation frequency, with values of 0.54, 0.52, and 0.66 for honey bee, bumble bee, and total pollinator visits, respectively. The BayesB model provided the most consistent prediction of fruit set, fruit weight, and seed set, with predictive abilities of 0.07, -0.08, and 0.42, respectively. Variable importance analysis revealed that genotypic differences in nectar volume had the greatest impact on honey bee and bumble bee recruitment, although preferences for flower morphological traits varied depending on the foraging task. Flower density was a major driving factor attracting nectar-foraging honey bees and bumble bees, while pollen-foraging bumble bees were most influenced by flower accessibility, specifically corolla length and the length-to-width ratio. Corolla length and the length-to-width ratio were also identified as the main predictors of fruit set, fruit weight, and seed count, suggesting that bumble bees and their foraging preferences may play a pivotal role in fruit production. Moderate to high narrow-sense heritability values (ranging from 0.30 to 0.77) were obtained for all floral traits, indicating that selective breeding efforts may enhance cultivar attractiveness to pollinators.

## INTRODUCTION

Southern highbush blueberry (*Vaccinium corymbosum* hybrids) relies on cross-pollination for optimal production, with bees contributing the vast majority of pollination (Taber and Olmstead, 2016; Mallinger et al., 2021a; Cortés-Rivas et al., 2023; Ramírez-Mejía et al., 2023). This places blueberry among the crops with the most demanding pollination requirements, as evidenced by substantial reductions in fruit set and berry quality in the absence of effective pollination (Benjamin and Winfree, 2014; Strik and Vance, 2019; Cavigliasso et al., 2021). To mitigate pollination-associated yield loss, U.S. blueberry producers spent more than $15M on beekeeping services, or $147.20 per acre in 2022 (USDA-NASS, 2023). Despite these substantial investments, blueberry still exhibits pollen limitation and subsequent yield loss (Sampson and Cane, 2000; Benjamin and Winfree, 2014; Campbell et al., 2018). As a result, recent reports have placed pollination and fruit set as a top priority for blueberry breeding, research, and development (Prasifka et al., 2018; Northwest Center for Small Fruits Research, 2020; personal communication, Florida Blueberry Extension Specialist Doug Phillips).

While traditionally treated as a management concern, blueberry pollination has shown inconsistent results in response to increasing honey bee hive stocking densities (Eaton and Nams, 2012; Arrington and DeVetter, 2018; Mallinger et al., 2021b). Moreover, honey bees have proven to be suboptimal pollinators of blueberry flowers due to their inability to "buzz" pollinate, which is essential to shed pollen from blueberry’s poricidal anthers (Javorek et al., 2002; Drummond, 2012; Cortés-Rivas et al., 2023). While bumble bees have been used to improve blueberry pollination outcomes, their stocking densities remain significantly lower than those employed for honey bees in most regions (Mallinger et al., 2021a). Consequently, efforts are underway to explore alternative solutions for improving pollination outcomes. Several studies have noted variability in pollinator attraction and fruit set among blueberry genotypes, prompting suggestions to breed pollinator-friendly cultivars with enhanced bee visitation, pollination efficacy, and crop yield (Lyrene, 1994; Prasifka et al., 2018).

Flower morphology and nectar rewards are among the key genetically controlled factors that contribute to pollinator attraction and pollination efficacy (Mitchell, 2004; Galliot et al., 2006; Heil, 2011; Escalante-Pérez and Heil, 2012; Quinet et al., 2016). Within this context, blueberry exhibits great variability for flower traits across species and genotypes (Eck and Mainland, 1971; Ritzinger and Lyrene, 1999; Sampson and Cane, 2000; Courcelles et al., 2013). Characterized by a unique bell-shaped corolla, blueberry flowers secrete nectar from the base of the flower, which is made accessible through a narrow aperture at the distal end of the flower. This structure impedes nectar access for some pollinators, such as honey bees, which often leads to nectar robbery and subsequent pollen limitation (Irwin et al., 2010). A previous study comparing four highbush blueberry (*V. corymbosum* L.) cultivars found that cv. ‘Duke’, the cultivar displaying the highest rates of honey bee visitation and fruit set, also had significantly wider corollas and aperture diameters than other cultivars, suggesting potential preferences for more accessible flowers (Courcelles et al., 2013). This study also identified an increased incidence of nectar robbery on the long and narrow flowers of cv. ‘Bluecrop’, which displayed significantly lower fruit set. Although distinct honey bee and bumble bee foraging patterns were noted between flower morphological traits, the limited scope of phenotypic diversity observed in this study was unable to resolve trait preferences between pollinator species.

Morphological traits can also influence fruit set through the promotion of outcrossing, which is crucial for circumventing fruit abortion due to self-incompatibility or early-acting inbreeding depression (Rick and Dempsey, 1969; Levy et al., 1978; Krebs and Hancock, 1990). Blueberry flowers exhibit herkogamy, wherein the stigma’s location within the corolla and the distance between the stigma and anthers reduce the incidence of self-pollination (Drummond and Rowland, 2020; Bieniasz and Konieczny, 2022). In wild lowbush blueberry *(V. angustifolium),* style length and the exertion of the stigma beyond the corolla were positively correlated with berry weight and the number of seeds per berry (Sampson et al., 2013), suggesting these traits may be beneficial for pollinator attraction or pollination success. While significant variation for herkogamy has been observed in highbush blueberry (Lyrene, 1994; Courcelles et al., 2013), its direct impact on seed and fruit set has yet to be examined.

Nectar volume and sugar concentration have shown to directly impact pollinator visitation in several crops, including blueberry, pepper, citrus, raspberry, blackberry, and sunflower (Jablonski et al., 1985; Rabinowitch et al., 1993; Albrigo et al., 2012; Schmidt et al., 2015; Mallinger and Prasifka, 2017). Early investigations found that nectar sugar concentration of lowbush blueberry (*V. angustifolium* and *V. myrtilloides*) was correlated with higher fruit weight for one of two species, although results were not congruent between years (Wood and Wood, 1963). Conversely, Jablonski et al. (1985) reported a positive association between nectar volume, pollinator visitation rates, and fruit set, but found no significant relationship with nectar sugar concentration. Importantly, this study found substantial diversity in nectar volume and sugar concentration across fourteen highbush blueberry cultivars. Therefore, it is suspected that the content of floral rewards may contribute to varietal differences in pollinator attraction and yield outcomes. However, preferences in nectar quantity and quality may vary between pollinator species. Additionally, the relative importance of nectar content with respect to other flower morphological traits has yet to be assessed.

Although several studies have found associations between nectar content and flower morphological traits with bee visitation or fruit set, current reports have focused on a small number of blueberry cultivars and wild species (Wood and Wood, 1963; Eck and Mainland, 1971; Ritzinger and Lyrene, 1999; Courcelles et al., 2013; Sampson et al., 2013). A comprehensive analysis that dissects the complex relationship between floral traits, bee behavior, and pollination outcomes is yet to be conducted for commercial southern highbush blueberries. Therefore, in the present study, we assessed the relationship between ten flower morphological and nectar traits, pollinator visitation, and three indices of pollination success, including fruit set, seed count, and berry weight, across 38 southern highbush blueberry genotypes, replicated over two years. Specifically, our objectives are three-fold: (i) assess the genotypic variability and heritability of floral traits within a diverse breeding population; (ii) test the importance of floral and nectar traits on pollinator attraction and multiple pollination and yield parameters; and (iii) determine the utility of these traits in prediction and selective breeding efforts.

## MATERIALS AND METHODS

### Plant Materials

This study included 38 southern highbush blueberry genotypes from the University of Florida Blueberry Breeding Program. These genotypes are part of a diverse breeding population comprising a total of 330 advanced-selected genotypes and were selected for their variability in fruit setting and overlapping flowering time based on data collected in 2019-2020. Plants were managed in a five-acre trial plot in Waldo, Florida, USA, under commercial management practices. Each genotype was clonally propagated with an average of 15 clones per genotype in the trial plot. Data was collected over two years during the 2021 and 2022 seasons, when plants were six and seven years of age, respectively.

### Pollinator management

A total of 60 honey bee hives were stocked adjacent to the experimental plot to serve the entire farm, according to the farm’s management practices using approximately 4 hives/acre. Bumble bee quads were also placed adjacent to the plot in both years with four quads being placed in 2021 and two quads in 2022 (0.8 quads/acre in 2021 and 0.4 quads/acre in 2022). In addition to managed bees, native insect pollinators could also have access to the plot throughout the bloom period.

### Flower morphological and nectar traits

Eight flower morphological traits were assessed, including corolla length, width, and length-to-width ratio, flower size (volume of cylinder, *V* ∼ πr^2^h), aperture diameter, style length, stigma protrusion from the corolla, and anther-stigma distance. During peak bloom, flower samples comprising ten recently opened flowers were collected from three random clones of each genotype, totaling 30 flowers per genotype each season. These flowers were cross-sectioned, imaged, and the traits measured via ImageJ FIJI software (Schindelin et al., 2012).

To evaluate nectar volume and sugar concentration, three randomly selected clones per genotype were covered with pollinator exclusion netting during the bud stage to prevent nectar consumption. During anthesis, 30 newly opened flowers were collected from each clone, and samples were promptly stored at 4°C until they could be processed within 24 hours in the laboratory. For each clone, the 30 flowers were pooled together, resulting in one composite sample per clone or three composite samples per genotype. Nectar volume was quantified using calibrated microcapillary pipettes (Drummond Scientific Co., Broomhall, Pennsylvania, USA), and nectar sugar concentration was measured using a hand-held refractometer adapted for small volumes (Bellingham and Stanley, College Station, Texas, USA).

### Pollinator visitation observations

Individual one-minute pollinator observations were conducted on each focal genotype to measure pollinator visitation and foraging habits. When weather conditions allowed, these observations were performed twice daily, at least three times per week, throughout the flowering period. Pollinator visitation was recorded as the number of flower visits for honey bees, bumble bees, and other flower visitors during each observation. Bumble bee foraging behaviors were also collected, discerning whether an individual was actively foraging for pollen or nectar. Pollen foraging was noted if an individual was carrying pollen in their corbiculae or if they were seen buzz pollinating. Those individuals that did not meet either of these criteria were determined to be nectar foraging. This distinction was not made for honey bees, which were nearly always foraging for nectar, or for other pollinators, which were few. In addition, the number of flowers on each bush, hereafter referred to as ‘flower density’, was also measured as the average number of observable flowers on a focal bush from a single vantage point during pollinator sampling.

### Fruit set and pollination indices

Various pollination indices, including seed count, fruit set, and fruit weight, were measured for three clones of each genotype. The proportion of fruit set was determined by dividing the number of ripe fruits by the number of flowers for three open-pollinated branches per clone. Fruit weight was recorded as the mass of 25 ripe fruits per clone, harvested during the peak window for each genotype. After weighing, ten berries were randomly selected for seed count measurements. Seed count comprised the total number of filled seeds, defined as large, round, and dark in color.

### Phenotypic correlation and heritability estimation

To determine the relationships between floral traits and pollination indices, a Pearson’s correlation test was conducted. Estimations of heritability were carried out via a linear mixed model, expressed in matrix notation as: 𝑦 = 1µ + 𝑋𝛽 + 𝑍𝑢 + 𝑒 (1), where ***y*** is the vector of phenotypic records for a specific trait; 𝝁 is the overall mean; 𝜷 is the fixed effect of year, ***u*** is the random effect of genetic values with 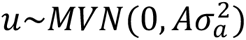 where ***A*** is the pedigree relationship matrix, 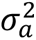 is the genetic variance, and ***e*** is the residual vector with 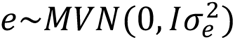, where ***I*** is the identity matrix and 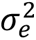 is the residual variance. ***X*** and ***Z*** are the incidence matrices for the fixed and random effects, respectively. Heritability was estimated using REML-BLUP methodology and is given by 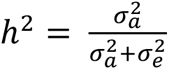. The pedigree relationship matrix was constructed using the ‘AGHmatrix’ R package (Amadeu et al., 2016), and mixed model analyses were conducted using the ‘ASReml-R’ package (VSNi, https://vsni.co.uk).

### Prediction of fruit set using pollinator visits

A linear mixed model was used to evaluate the predictive ability of pollinator visitation rates for fruit set across genotypes. This model treated pollination visits as a fixed effect and included both a random intercept and a random slope for pollination visits, enabling the prediction of each genotype’s independent response to pollinator visitation. The random regression was conducted using the ‘GMMAT’ R package (Chen et al., 2023).

### Prediction of pollinator visitation and pollination indices using flower traits

We tested four statistical and machine learning model, including partial least squares (PLS), ridge-regression (RR), BayesB, and random forest (RF), for their ability to predict pollinator visitation frequency and indices of pollination success based on flower traits. Predictive abilities were assessed through a k-fold cross-validation process, using training set (80%) to test set (20%) ratio repeated 100 times. For statistical methods, we employed the linear model 𝒚 = 𝟏µ + 𝐗𝛃 + 𝐖𝐬 + 𝐞 (2), where ***𝐲*** is the vector of phenotypic records for a particular trait; ***𝜇*** is the overall mean; 𝜷 is the fixed effect of year with incidence matrix ***X****; **W*** is the matrix containing the flower traits, ***s*** is the vector of their effects on the pollinator visitation and pollination indices, and ***e*** is the residual vector with 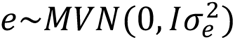, where ***I*** is the identity matrix and 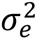 is the residual variance.

Each statistical methodology comes with its own set of assumptions: PLS assumes that all effects within the model (2) are fixed effects and employs latent variables and the ordinary least squares method for estimating these effects. RR assumes ***s*** as random effects following a normal distribution with a common variance for all effects and utilizes REML-BLUP methodology for estimating these effects. BayesB is a Bayesian method and assumes ***s*** as random effects following a mixture of normal distribution with a proportion of effects that can be null effects, determined by their importance. As a representative of the machine learning approach, we used random forest (RF), a method based on tree algorithms that uses a third of the number of variables randomly sampled as candidates at each split (Hastie et al., 2009). The specific implementations include: PLS was fitted via the ‘pls’ R package (Mevik and Wehrens, 2007), BayesB and RR were fitted via the ‘BGLR’ R package (Pérez and De Los Campos, 2014) and ‘rrBLUP’ R package (Endelman, 2011), respectively. RF was fitted via the ‘randomForest’ R package (Liaw and Wiener, 2002). In the Bayesian method, we used 300,000 total iterations for the Markov chain Monte Carlo algorithms and the first 20,000 iterations were discarded as burn-in. After every set of 5 iterations (thin), a sample was retained to calculate posterior estimates. The convergence of the Markov chains was verified through Geweke’s diagnostic (Geweke, 1992).

### Variable importance analysis

Based on the results of previous prediction analyses, we selected the model that demonstrated superior performance in predicting pollinator visitation and pollination indices. Variable importance was determined by calculating the relative influence of each variable in relation to these specific pollination traits. For statistical methods, we considered the value of the normalized regression coefficient, while for the random forest method, we utilized a measure based on the usage of a specific variable at each split in each tree. The loss in the split-RSME (Root-Mean-Square Error) is the importance measure attributed to the splitting variable and is accumulated over all of the trees in the forest individually for each variable. To enable meaningful comparisons across traits, all variance importance values were standardized.

## RESULTS

### Phenotypic diversity, correlation, and heritability

Considerable phenotypic variability was observed for all flower traits across the 38 southern highbush blueberry genotypes (Figure 1; Supplementary Figure S1).

**Figure 1.**
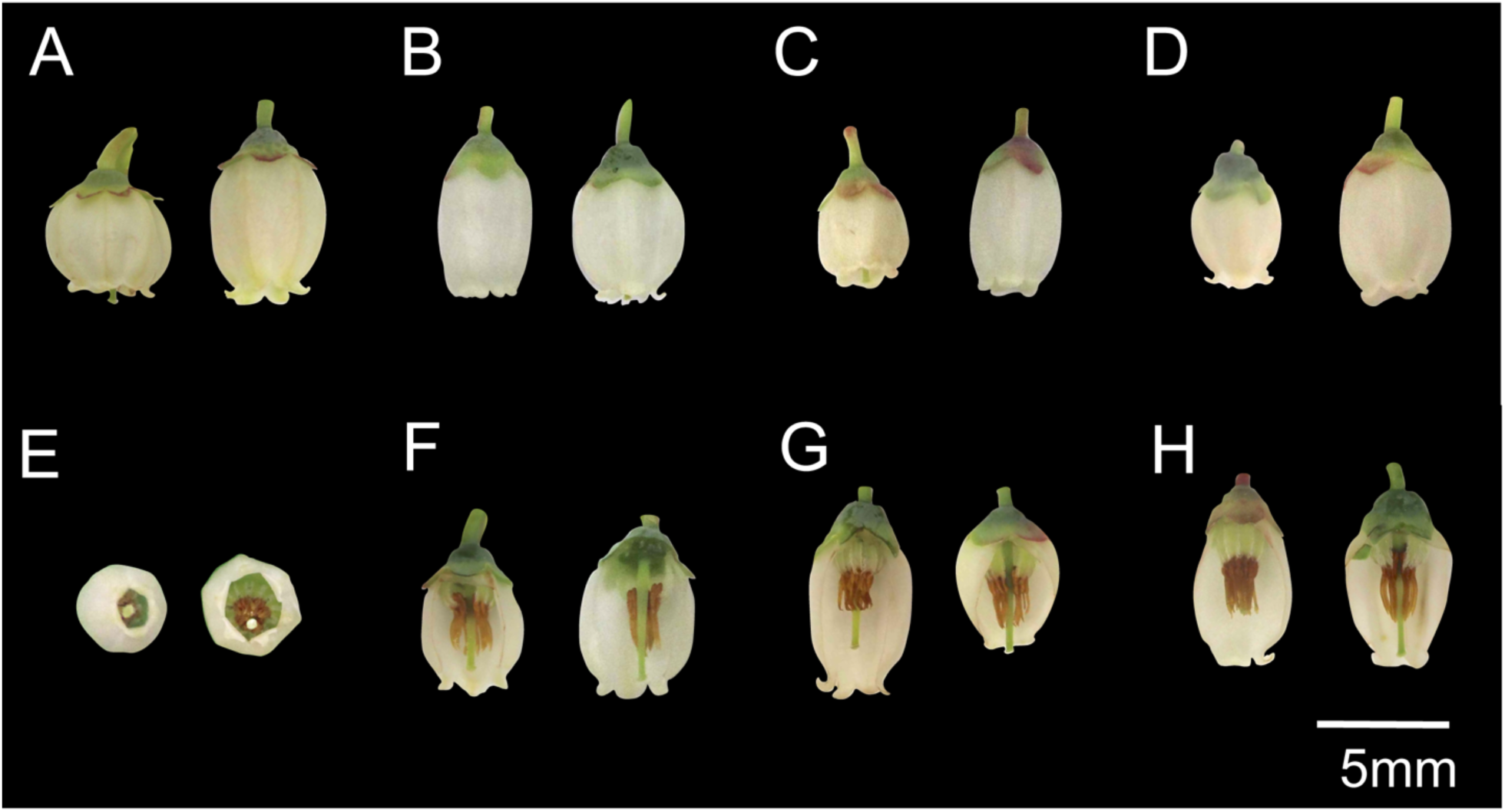
Floral diversity observed between 38 southern highbush blueberry genotypes for (A) corolla length, (B) corolla width, (C) ratio of corolla length-to-width, (D) flower size, (E) aperture diameter, (F) style length, (G) stigma protrusion, and (H) anther-to-stigma distance.

A Pearson’s correlation test revealed significant associations between flower size and nearly all morphological traits, except for corolla aperture diameter and the corolla length-to-width ratio (P < 0.05; Figure 2). Nectar volume and sugar concentration were negatively correlated (*r* = -0.38, P < 0.001), and the volume of nectar secreted was positively associated with flower size (*r* = 0.31, P < 0.001). A significant correlation between the pollination indices of seed count and fruit weight was also observed (*r* = 0.22, P < 0.01). The heritability values of all flower morphology and nectar traits were moderate to high, ranging from 0.30-0.77 (Figure 2). Pollination indices showed moderate heritability values of 0.31, 0.53, and 0.42 for fruit set, seed count, and fruit weight, respectively.

**Figure 2.**
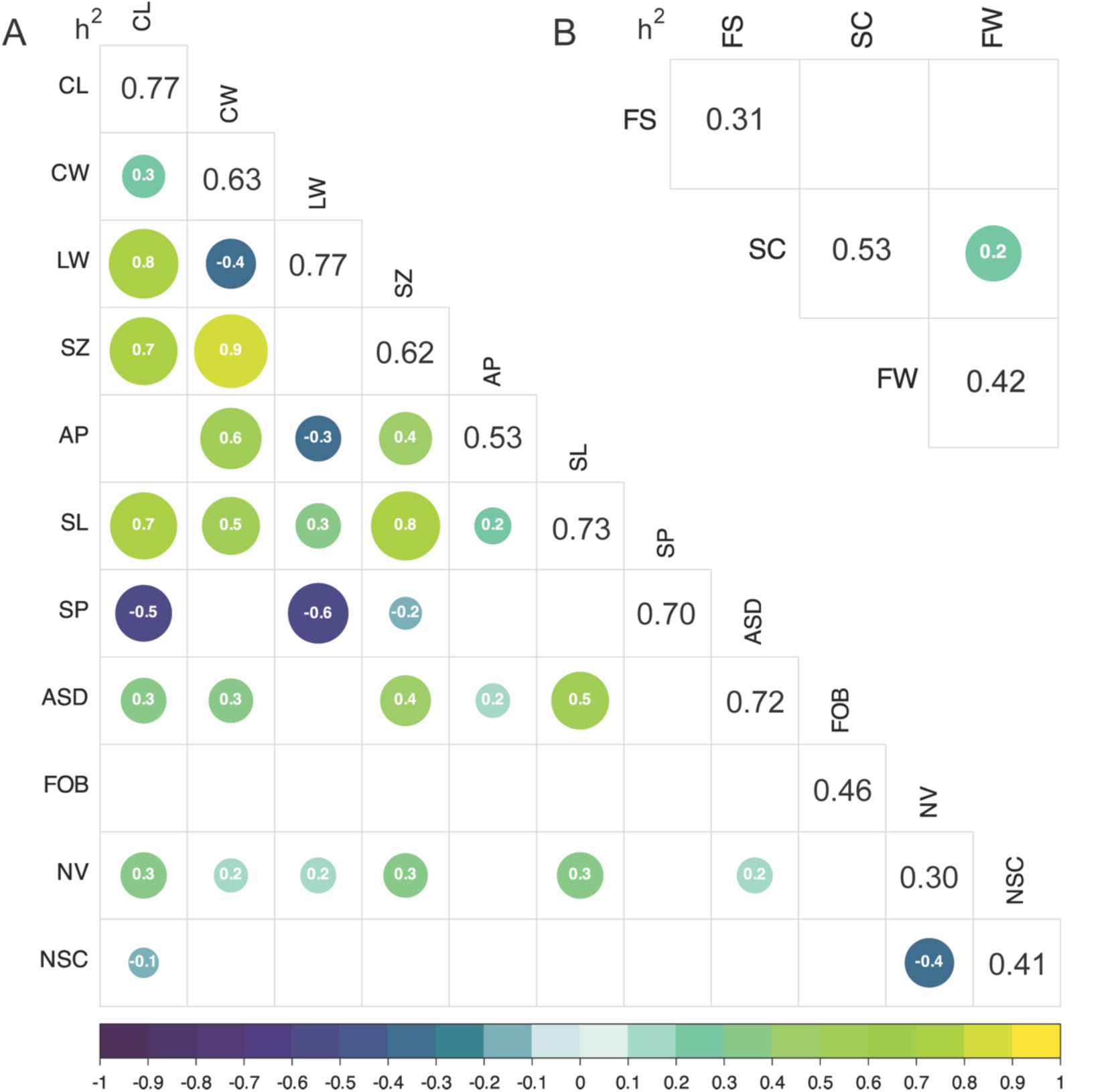
Narrow-sense heritability (diagonal) and Pearson’s correlation coefficients (off-diagonal) of (A) flower traits, and (B) pollination indices. The size and color of each dot is proportional to the magnitude and direction of the Pearson’s correlation coefficient, respectively. Only statistically significant relationships with *p* < 0.05 are displayed. CL, corolla length; CW, corolla width; LW, ratio of corolla length-to-width; SZ, flower size; AP, aperture diameter; SL, style length; SP, stigma protrusion from corolla; ASD, anther-to-stigma distance; FOB, flowers on bush; NV, nectar volume; NSC, nectar sugar concentration; FS, fruit set; SC, seed count; FW, fruit weight.

### Genotypic differences in pollinator visitation

The total pollinator visits observed for each genotype ranged between 30 to 218 and 0 to 130 in the 2021 and 2022 seasons, respectively (Figure 3; Supplementary Figure S2). In the 2022 season, there was a decrease in visitation rates across all genotypes. This decline could be due to the earlier bloom period observed in 2022 or variability in temperature and cloud cover, which are known to impact bee activity (Supplementary Figure S3). Most pollinator visits (90%) were made by honey bees (*Apis mellifera*), which exclusively foraged for nectar. Bumble bees, including *Bombus impatiens* and *Bombus bimaculatus*, comprised 8% of flower visits, of which 75% foraged for nectar, and the remaining 25% collected pollen. Other species comprised 2% of total flower visitors and consisted mostly of carpenter bees (*Xylocopa virginica* and *X. micans*), although a small number of visits were also made by southeastern blueberry bees (*Habropoda laboriosa*), flower wasps (*Scoliid spp*.), and hover flies (Syrphidae family).

**Figure 3.**
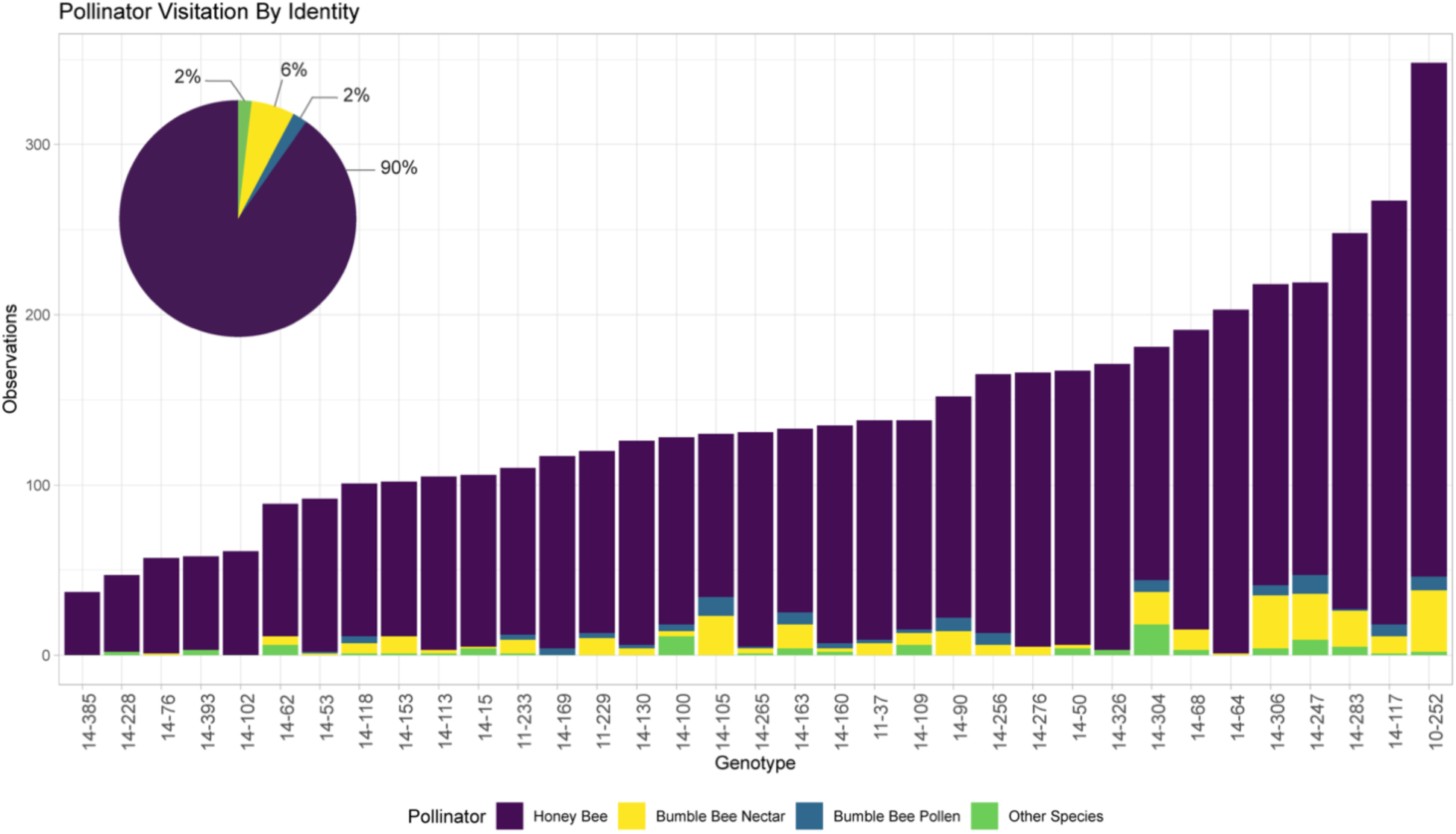
Total pollinator visitation (pie chart) and pollinator visitation per genotype (bar plot), represented as the total number of pollinator visits across both the 2021 and 2022 season. Colors indicate pollinator species and foraging behavior. Other species included carpenter bees, flower wasps, hover flies, and the southeastern blueberry bee.

In certain genotypes, honey bees comprised only 73% of all visits, while others were solely visited by honey bees. Additionally, we noted distinct preferences among bumble bees for specific genotypes, with their representation reaching up to 26% of all visits, while some genotypes received zero visits from bumble bees. Visits from other pollinator species were generally infrequent, and never exceeded 10% of the total pollinator visits for any genotype.

### The effect of pollinator visitation on fruit set was not significant across genotypes

A linear mixed model was used to predict fruit set using pollinator visits in 38 genotypes. However, the genetic effect was statistically non-significant when we evaluated the performance of the genotypes individually. These genotypes demonstrated vast differences in fruit set regardless of visitation, suggesting that genotypic variability for fruit set may be mediated by factors other than pollinator visitation. Additionally, the relationship between pollinator visitation and fruit set varied between years (Figure 4). In 2021, all genotypes displayed higher fruit set with increasing pollinator visitation frequency, but this pattern was not observed in 2022.

**Figure 4.**
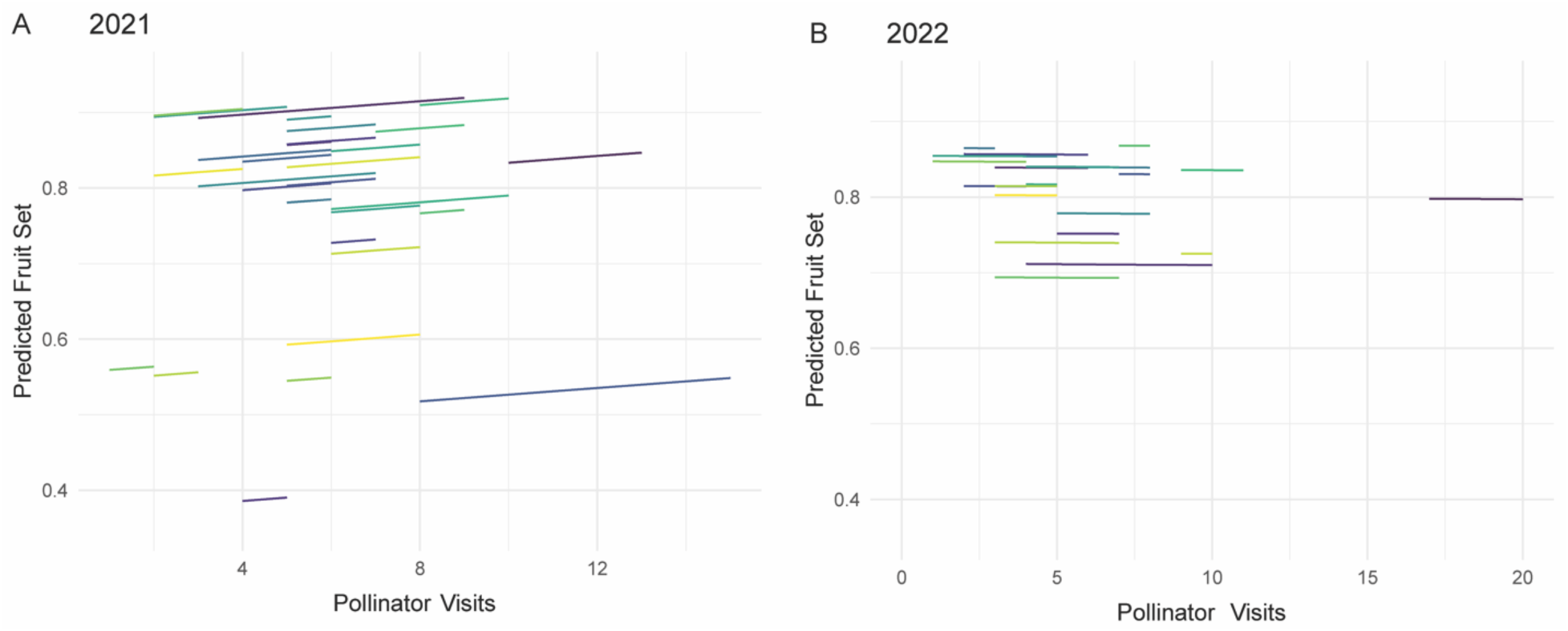
Fruit set predictions based on total pollination visits normalized by the number of flowers per bush during the 2021 and 2022 seasons. Each line represents a unique genotype.

### Prediction models and variable importance analyses indicated floral trait preferences between bee species

PLS, BayesB, RR, and RF regression models were evaluated for their ability to predict pollinator visitation frequency and pollination indices using flower density, nectar rewards, and flower morphological traits as predictor variables (Figure 5; Supplementary Figure S6). To determine if spatial variation contributed to pollinator visitation or pollination indices, row and column were fitted as fixed effects. Neither row nor column were significant in both years, hence, they were excluded from the final model (Supplementary Figure S4). Additionally, three inconsistent genotypes were removed from the analysis since they likely maintain a high degree of self-compatibility and displayed high seed numbers despite low pollinator visitation rates (Supplementary Figure S5).

**Figure 5.**
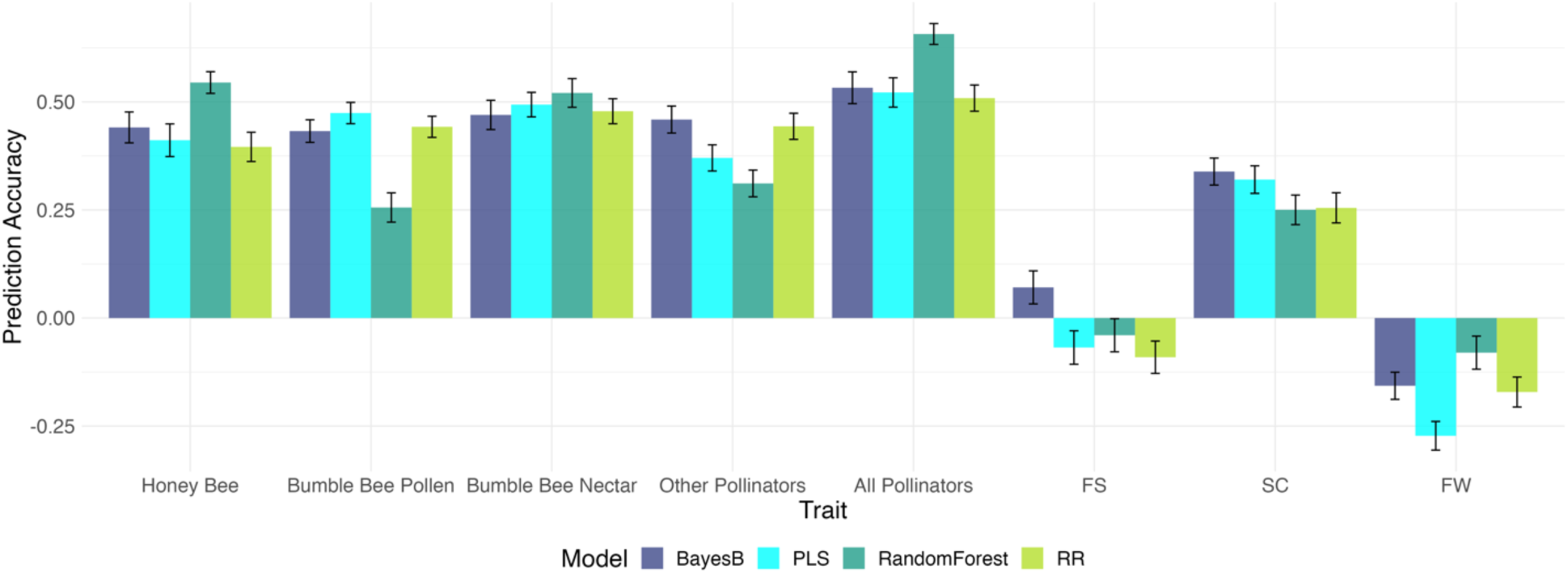
Predictive abilities for pollinator visitation and pollination indices using floral traits. Pollination indices include fruit set (FS), fruit weight (FW), and seed count (SC). Four prediction models were tested: BayesB, partial least squares (PLS), random forest, and ridge-regression (RR) models.

RF exhibited the highest predictive abilities for honey bee, nectar foraging bumble bee, and overall pollinator visitation frequency, with predictive abilities of 0.54, 0.52, and 0.66, respectively (Figure 5). Therefore, RF models were employed to explore the relative contribution of each floral trait to the prediction of pollinator visitation in the variable importance analysis. Although the predictive abilities for pollination indices were generally low across all models, BayesB performed best, with values of 0.07, 0.34, and -0.16 for fruit set, seed count, and fruit weight, respectively.

### The influence of floral traits on pollinator visitation rates depended on foraging task

The relevance of each floral trait in the prediction of pollinator visitation frequency and pollination success is shown in Figure 6 and Supplementary Figure 6. The analysis of variable importance uncovered variations in trait preferences among different bee species (Figure 6A). Particularly, nectar volume emerged as the most influential predictor of honey bee visitation, exerting a stronger effect on honey bees compared to bumble bees and other species. According to the partial dependence profile derived from random forest regression models, honeybee visitation increased with larger volumes of nectar secretion, reaching a plateau near 400 ml per 30 flowers (∼13ml/flower) (Supplementary Figure S6D). Additionally, honey bee visits were strongly impacted by the diameter of the corolla aperture compared to other pollinator species.

**Figure 6.**
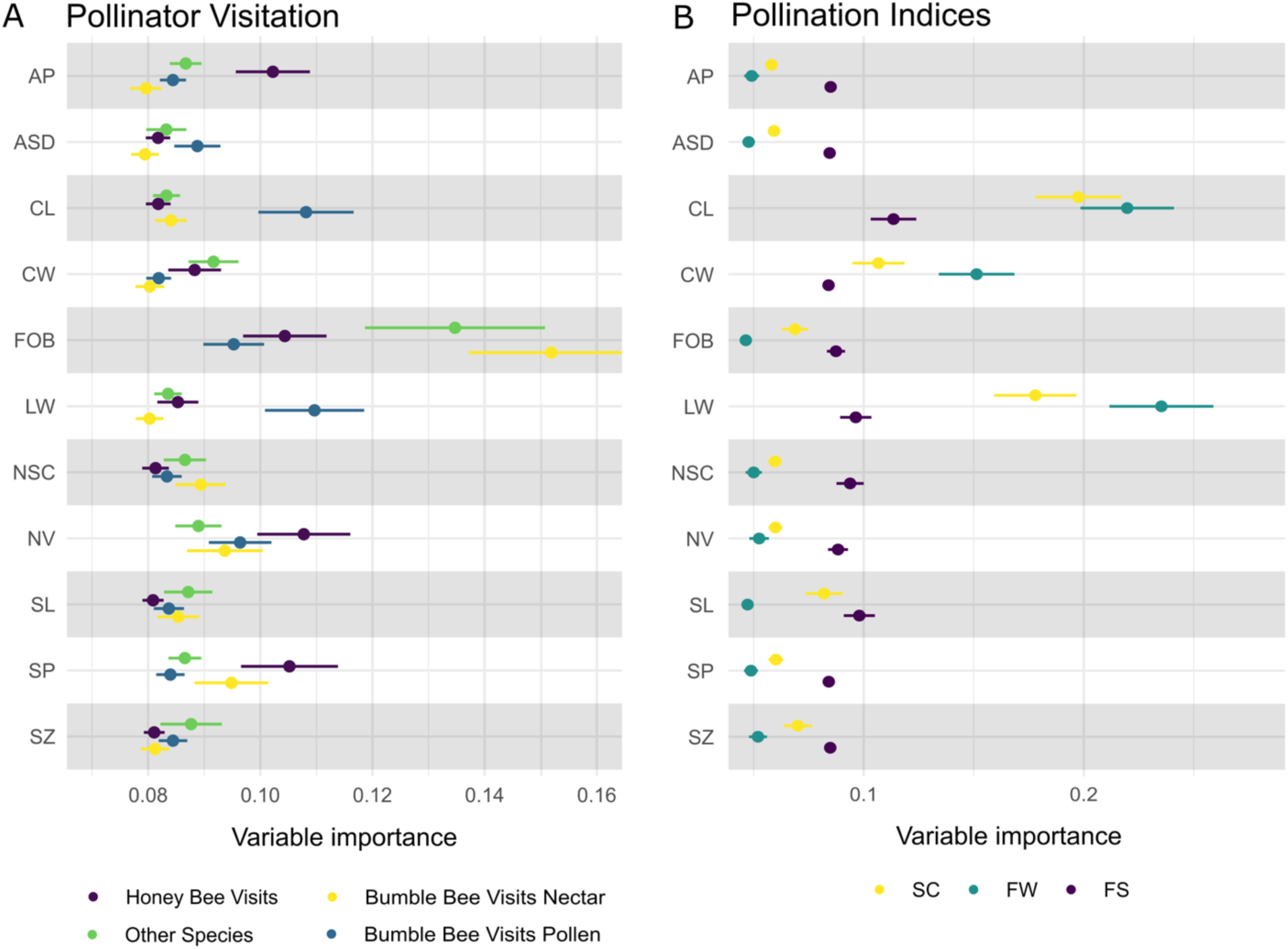
Variable importance shows the relevance of specific flower traits in predicting pollinator visitation (A) and pollination indices (B), using random forest and BayesB models, respectively. Higher values indicate greater importance. Flower traits: AP, aperture diameter; ASD, anther-to-stigma distance; CL, corolla length; CW, corolla width; FOB, flowers on bush (flowering density); LW, ratio of corolla length-to-width; NSC, nectar sugar content; NV, nectar volume; SL, style length; SP, stigma protrusion from corolla; SZ, flower size. Pollination indices: SC, seed count; FW, fruit weight; FS, fruit set.

Nectar-foraging insects, encompassing honey bees, bumble bees, and other species, displayed marked associations with flower density (Figure 6A). In contrast, pollen-foraging bumble bees displayed minimal responsiveness to flower density and were instead influenced by accessibility traits, namely corolla length and the corolla length-to-width ratio. Genotypes with protruded stigmas displayed higher visitation rates from honey bees and nectar-foraging bumble bees. However, this trait did not significantly contribute to the prediction of fruit set, fruit weight, or seed count (Figure 6B).

## DISCUSSION

Since the early stages of blueberry domestication, there has been extensive documentation of how pollination benefits fruit set and yield (Shaw et al., 1939; Eck and Mainland, 1971; Marucci and Moulter, 1977; Lyrene, 1990; Williamson and Nesmith, 2007). Despite the routine use of pollination services, there is considerable variation in pollination success and subsequent yield reported among blueberry cultivars (Marucci, 1966; Ehlenfeldt, 2001; Taber and Olmstead, 2016). Flower morphological and nectar-related traits have shown to play significant roles in pollinator attraction, pollination efficacy, and fruit production (Eck and Mainland, 1971; Ritzinger and Lyrene, 1999; Courcelles et al., 2013). Selective breeding of these traits has been proposed to address pollination deficits and improve yield outcomes (Lyrene, 1994; Prasifka et al., 2018). However, prior studies have been predominantly centered on northern highbush cultivars or a small selection of wild species and inter-specific hybrids (Eck and Mainland, 1971; Lyrene, 1994; Ritzinger and Lyrene, 1999; Courcelles et al., 2013; Sampson et al., 2013). Therefore, our understanding of genotypic diversity and the heritability of these traits in southern highbush blueberry remains limited, even though such knowledge is crucial for breeding efforts. In the present study, we assessed the diversity of flower morphology and nectar-related traits in 38 diverse southern highbush blueberry genotypes and evaluated their relative importance for pollinator attraction and pollination success within a breeding context.

The genotypes included in this study exhibited considerable diversity in all flower and nectar traits. Additionally, we found all traits to be moderately to highly heritable (0.30-0.77), which suggested that relevant genetic progress can be achieved for these traits throughout breeding cycles. This observation concurs with Lyrene (1994), which reported that environmental conditions, including chilling units and temperature, had little impact on flower morphology, suggesting strong genetic control of floral traits. Nectar volume exhibited the lowest heritability value (h^2^ = 0.30). Nectar secretion is known to fluctuate with factors such as flower age, time of day, and cultivar, which could affect the estimation of genetic parameters (Wood and Wood, 1963; Mitchell, 2004; Nepi and Stpiczyńska, 2008).

Honey bees displayed a clear preference for genotypes with larger nectar volumes, which is consistent with previous studies that demonstrated a positive correlation between nectar volume and bee visitation across various blueberry cultivars (Wood and Wood, 1963; Brewer and Dobson, 1969; Jablonski et al., 1985). Our study also found a negative association between nectar volume and sugar concentration, indicating a dilution effect in nectar secretion. Distinct preferences were also apparent based on foraging tasks, with nectar-foraging bumble bees favoring genotypes with higher flower density and nectar volume. In contrast, pollen-foraging bumble bees showed a preference for flowers with short and wide corollas, facilitating stamen access. As the long tongue of bumble bees enables access to nectar from a broad range of flower shapes and sizes (Goulson, 2010), varietal preferences may be primarily constrained by flower density and not individual blueberry flower shape and size.

Honey bees have previously been shown to prefer larger blueberry flowers with wide apertures (Courcelles et al., 2013). To our surprise, we observed little to no relationship between flower size and honey bee visitation. Interestingly, random forest regression captured a non-linear relationship between honey bee visitation and aperture diameter.

Honey bee visits were highest for genotypes with small aperture diameters, narrower than 3.5mm. However, once apertures exceeded a threshold of ∼3.5 mm, honey bee visitation began to increase. While it is still unclear why we observed this pattern, one possibility is that the narrow opening may restrict competing larger-bodied bees, such as carpenter bees and bumble bees, allowing for exclusive access to honey bees. Honey bees and bumble bees have been shown to compete for nectar, especially when hives are placed near bumble bee quads (Thomson, 2004). These competitive effects lead to altered foraging behavior and often decreased nectar availability (Page and Williams, 2023). In this case honey bees could be favoring flowers with smaller apertures that reduce competition from bumble bees.

A key expectation of this work was that genotypes exhibiting greater pollinator visitation frequency would also exhibit higher fruit set. However, we did not find this relationship to be statistically significant. The constrained variability observed for fruit set could have limited our capacity to establish this connection. Additionally, factors outside of pollen load may affect fruit set, including environmental conditions during the bloom period, plant health, and fertility (Davies, 1986; Yang et al., 2019; Drummond, 2020). Still, our analyses showed that each genotype benefited from increased pollinator visitation frequency. Moreover, the variable importance analysis revealed similar trends in the traits contributing to the predictive ability of both pollinator visitation and fruit set, suggesting an indirect relationship.

Corolla length and width were identified as the main drivers of pollen-foraging bumblebee visitation and all indices of pollination success, including fruit set, seed count, and fruit weight. This finding suggests a direct connection between flower morphology, pollinator recruitment, and fruit production. Previous research has demonstrated that pollen-foraging bumble bees are nearly five times more effective at pollen transfer than honey bees (Javorek et al., 2002). Therefore, it is possible that, despite the lower frequency of visits made by pollen-foraging bumble bees, they had a disproportionate effect on pollination indices due to their higher pollination efficacy. Furthermore, corolla length and the corolla length-to-width ratio had the highest heritability among all traits (h^2^ = 0.77), indicating that selective breeding for these traits may improve pollination outcomes at a higher rate than other flower characteristics.

The observed variation in the relative importance of blueberry floral traits across pollinator species and foraging behaviors is a novel finding with great implications for future efforts to enhance pollination. The strong correlation and high heritability of these traits further emphasizes the potential of plant breeding to improve pollination outcomes and benefit growers through higher fruit production.

## CONCLUSIONS

This study delved into the multifaceted relationship between pollinators, flower morphology, pollination success, and yield parameters in blueberry, a subject that has been under-explored in the context of plant breeding. Our analyses confirmed the importance of nectar reward content and flower morphological traits in pollinator attraction. Moreover, we found these traits to be highly heritable, providing justification for breeding to improve pollination outcomes. Random forest and BayesB regression resulted in moderate to high predictive abilities for bee visitation and pollination success using floral traits. Subsequent variable important analysis revealed distinct preferences between honey bees and bumble bees, as well as between bees conducting different foraging tasks. Nectar volume was found to be the most important trait mediating honey bee visitation frequency, while bumble bees were instead influenced by flower accessibility traits, including corolla length and the length-to-width ratio. Flower density had a substantial impact on all pollinator species, with a pronounced effect on nectar-foraging bumble bees and non-honey bee species. These findings emphasize the critical role of floral reward density and flower accessibility in the pollination landscape. While the relationship between flower visitation frequency and fruit set was not significant in our analyses, we expect that using more pollinator-attractive cultivars on farms experiencing pollination deficits may result in higher production, given the generally positive effect of pollination on yield. Additionally, the high heritability for floral traits reported here can provide valuable insight for the prioritization of traits for the genetic improvement of pollinator attraction, and ultimately yield in an outcrossing and pollinator-dependent crop such as blueberry.

## ACKNOWLEDGEMENTS

This study was supported by royalties from the University of Florida Blueberry Breeding Program and funding from Southern SARE (Graduate Student Grant – GS21-244). The authors are grateful to Straughn Farms LLC for enabling research experiments to be conducted on their farm and for continued support of the University of Florida Blueberry Breeding program. We also thank Dr. Ivone de Bem Oliveira for important discussions that helped develop the project idea, Dr. Felix Enciso-Rodriguez, and Dr. Juliana Benevenuto for reviewing the final manuscript, and Mia Acker, Dorothy Coutu, Ben Covert, Ryan Cullen, Haley Dabbs, Jordyn Lind, and Taylor Sawyer, and who assisted in sample processing and phenotyping efforts.

## AUTHOR CONTRIBUTIONS

JC and PRM conceived the study. JC designed the experiment and population for the study. JC and AK conducted the phenotyping of floral traits and pollination indices. JJT conducted pollinator observations. JC, PA, and CA carried out statistical analyses. JC wrote the manuscript with contributions from all authors.

## DATA AVAILABILITY STATEMENT

Data will be made available upon request.

## SUPPLEMENTARY MATERIALS

**Figure S1.**
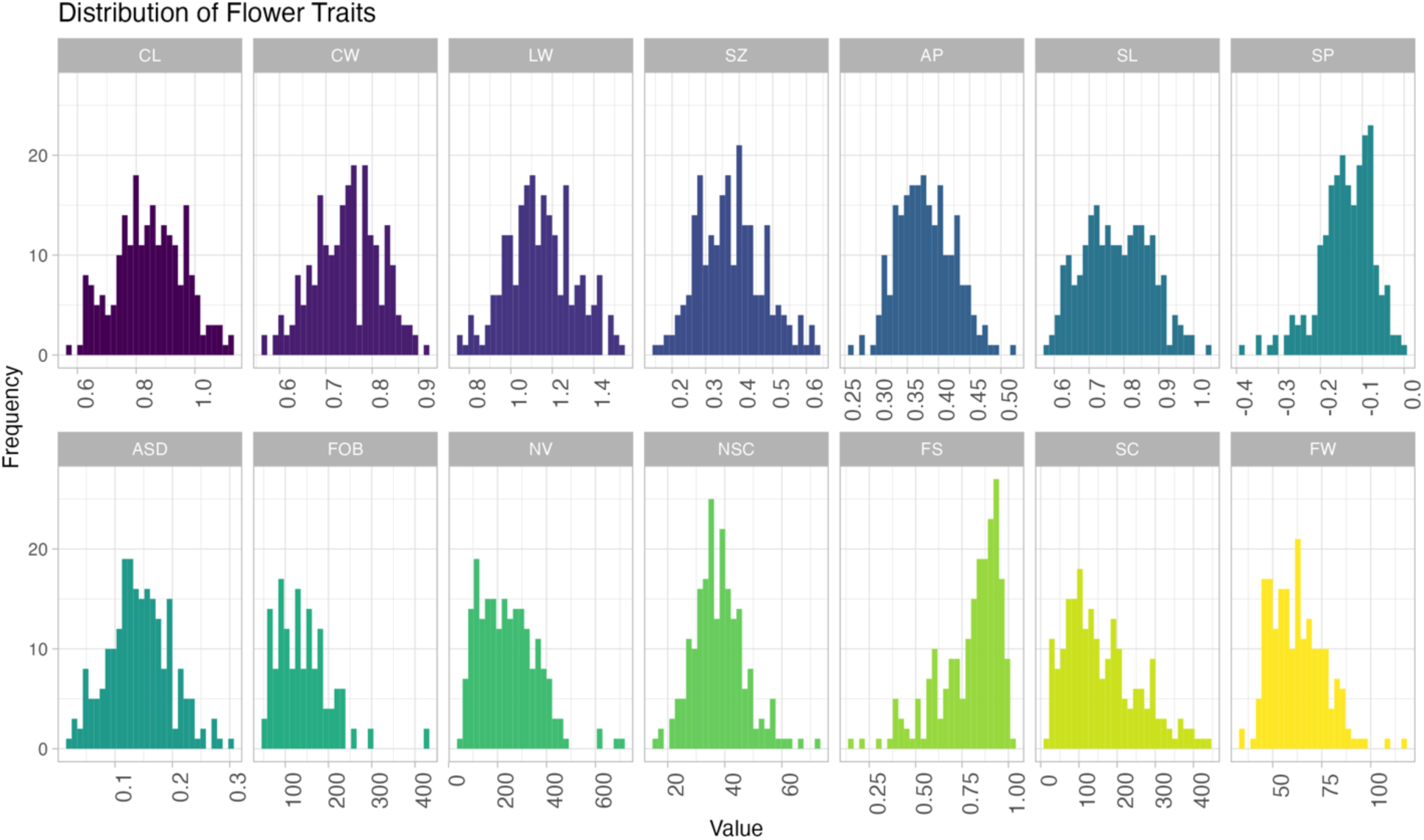
Histogram distribution of flower morphological traits. CL, corolla length (cm); CW, corolla width (cm); LW, ratio of corolla length-to-width; SZ, flower size (cm^3); AP, aperture diameter (cm); SL, style length (cm); SP, stigma protrusion from corolla (cm); ASD, anther-to-stigma distance (cm); FOB, flowers on bush (flowering density); NV, nectar volume (µL); NSC, nectar sugar content (°BRIX); FS, fruit set; SC, seed count of 10 berries; FW, fruit weight of 25 berries (g).

**Figure S2.**
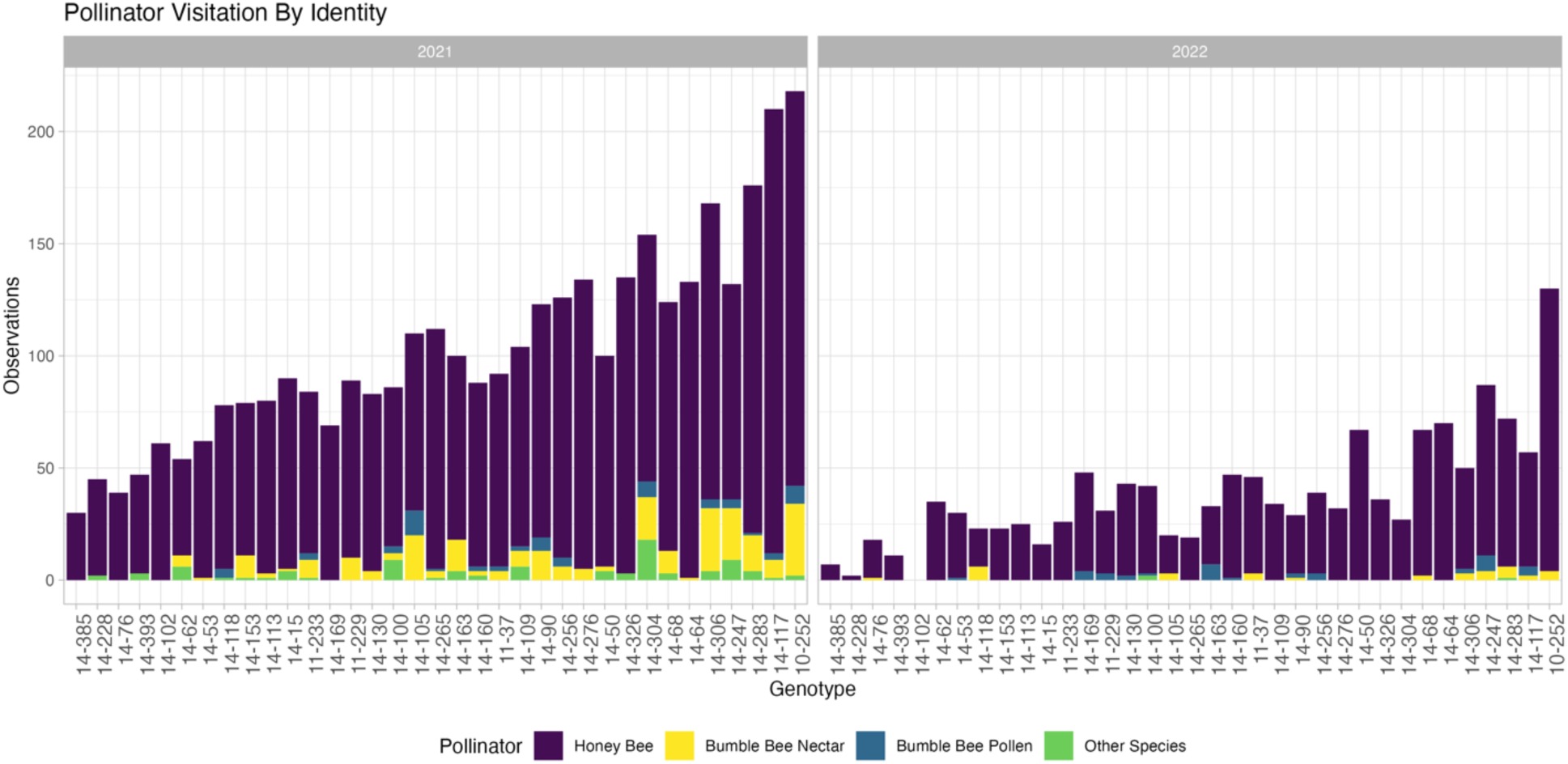
Pollinator visitation for each genotype between years. Colors indicate pollinator species and foraging behavior. Other species included carpenter bees (*Xylocopa virginica* and *X. micans*), flower wasps (*Scoliid* spp.), hover flies (Syphridae), and the southeastern blueberry bee (*H. laboriosa*).

**Figure S3.**
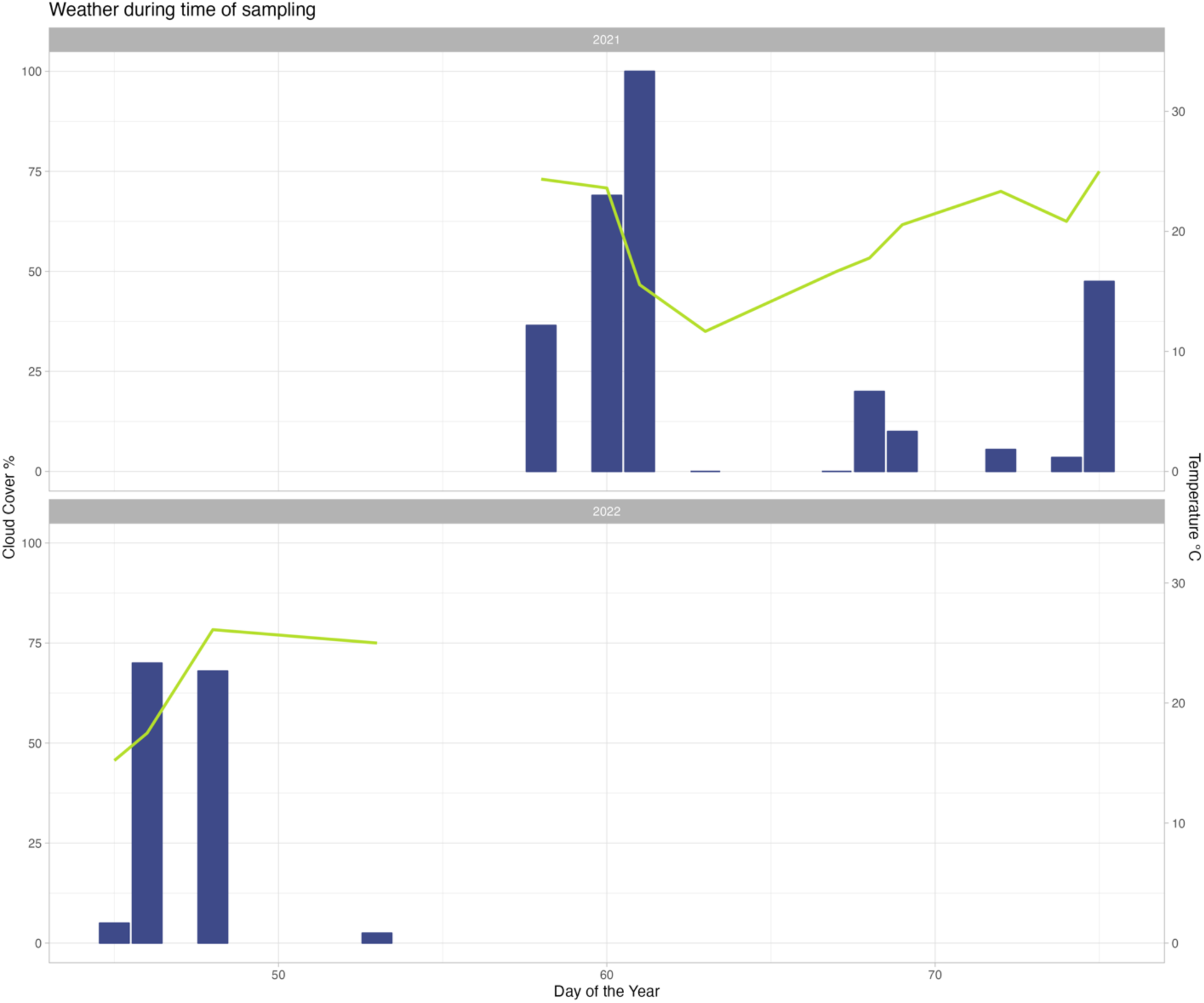
The temperature (°C) (green line) and cloud-cover percent during the 2021 and 2022 growing seasons. Julian date is presented as day of the year.

**Figure S4.**
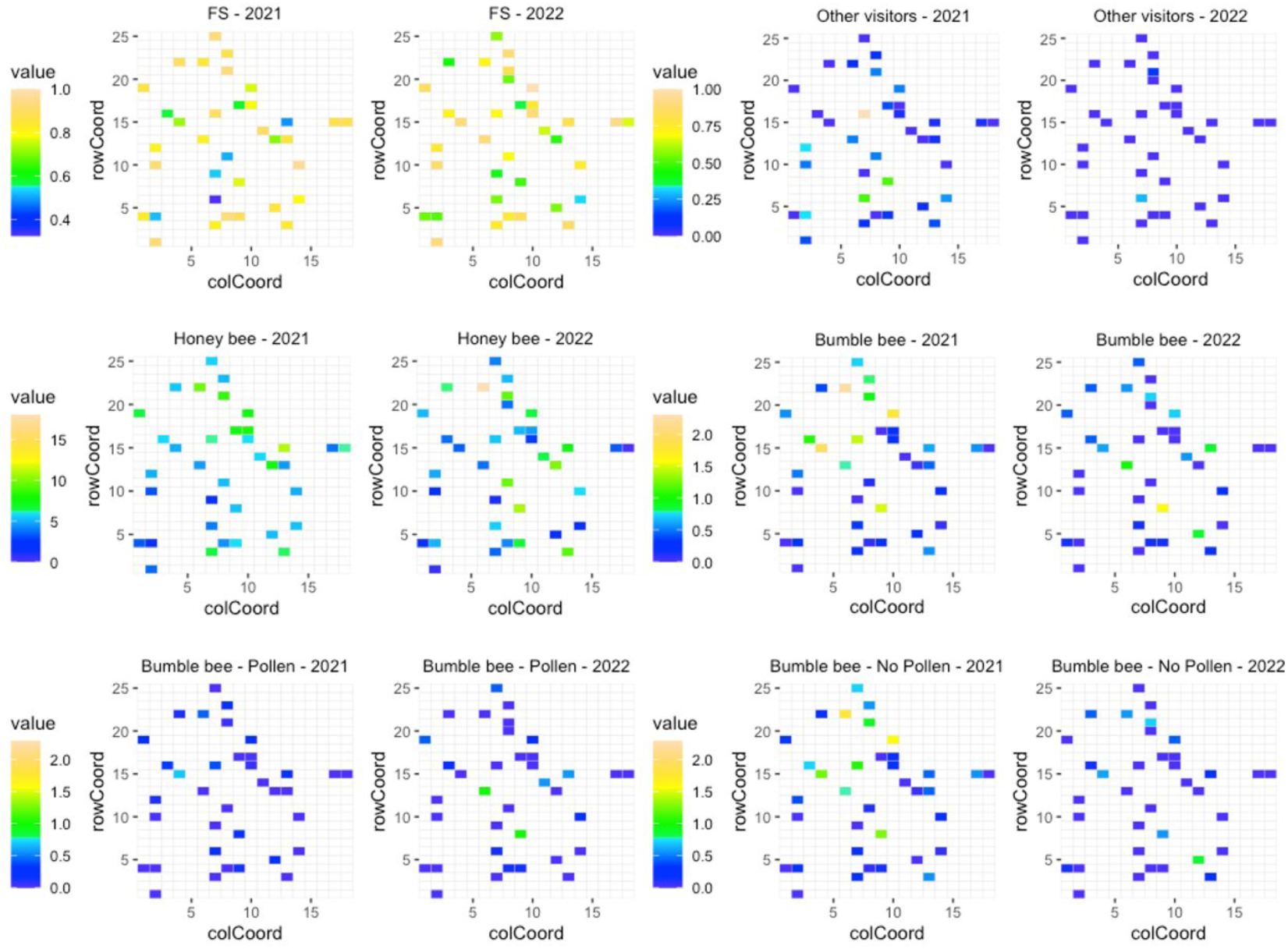
The spatial distribution of fruit set (FS), honeybee visitation, bumblebee visitation, and other flower visitors across row and column positions in the 2021 and 2022 seasons.

**Figure S5.**
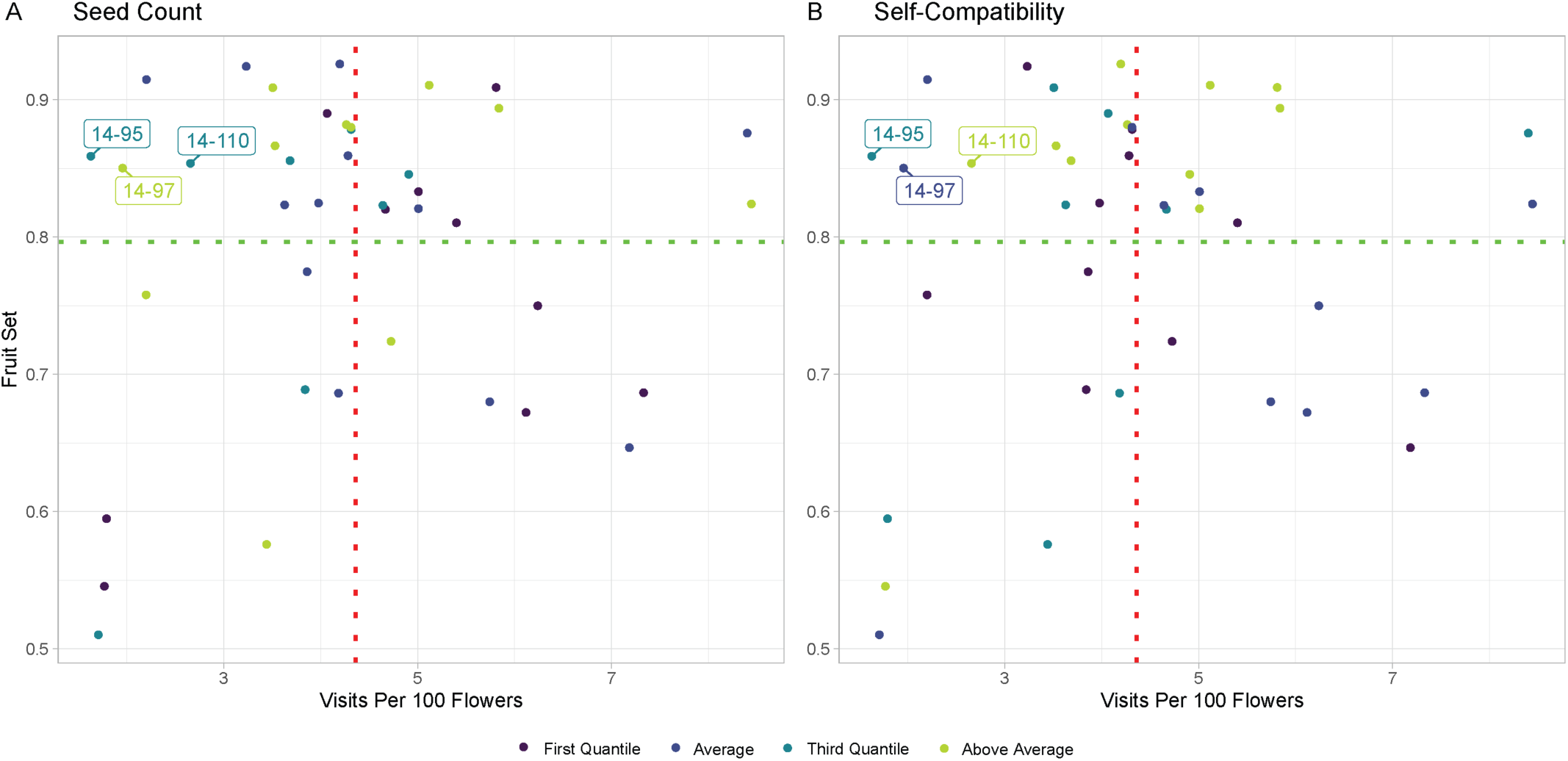
Outliers removed due to high number of seeds with low pollinator observations (A) and high rates of self-compatibility (B), inflating fruit set at low visitation rates.

**Figure S6.**
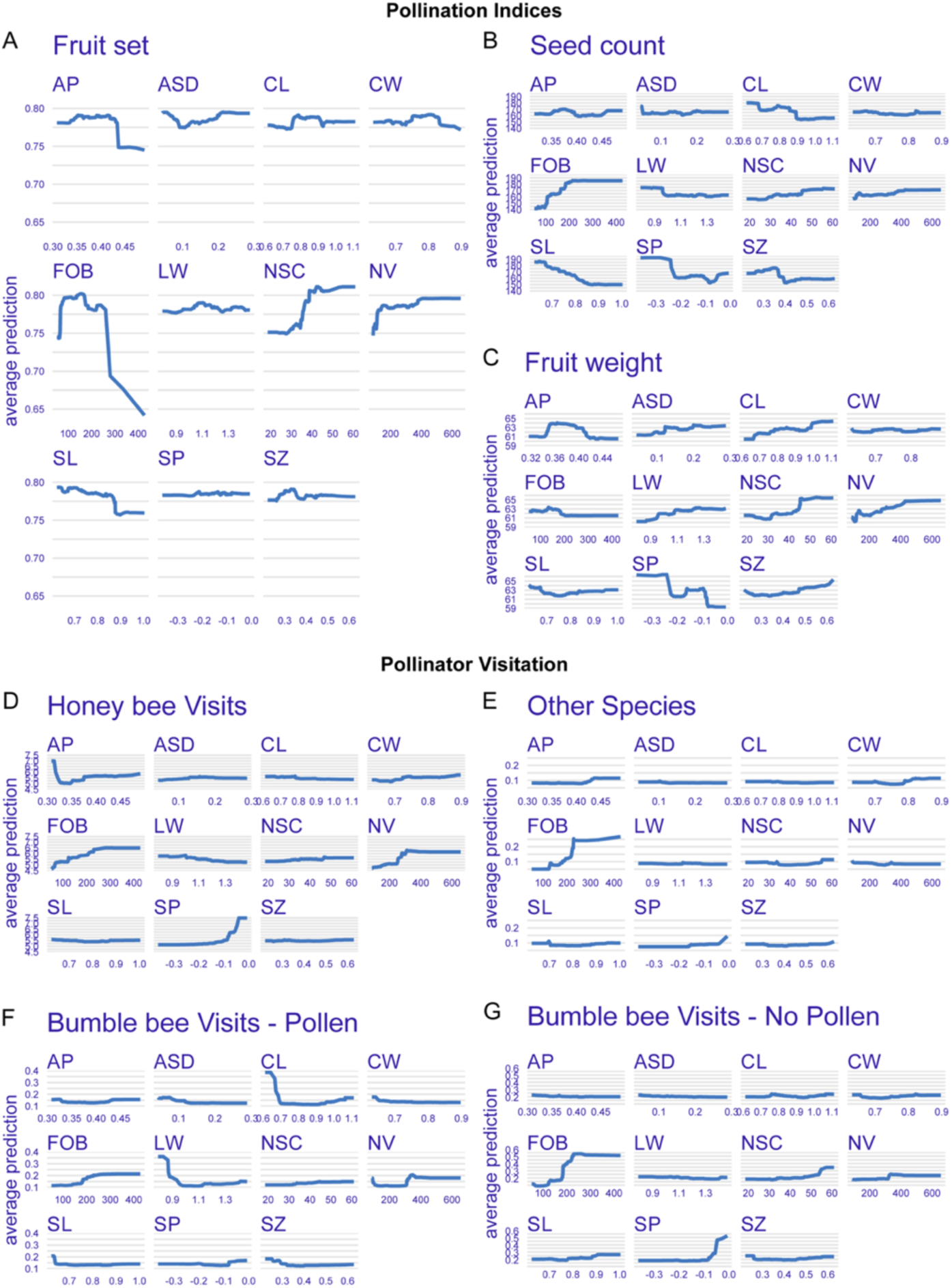
The partial dependence profile for the relationship between pollination indices (A-C) and pollinator visitation frequency (D-G) with each individual flower trait based on BayesB and Random Forest regression, respectively. AP, aperture diameter; ASD, anther-to-stigma distance; CL, corolla length; CW, corolla width; FOB, flowers on bush (flowering density); LW, ratio of corolla length-to-width; NSC, nectar sugar content; NV, nectar volume; SL, style length; SP, stigma protrusion from corolla; SZ, flower size.

