## Supplementary Material for "Genotypic variation in blueberry flower morphology and nectar reward content affects pollinator attraction in a diverse breeding population"

**Supplementary Figures**

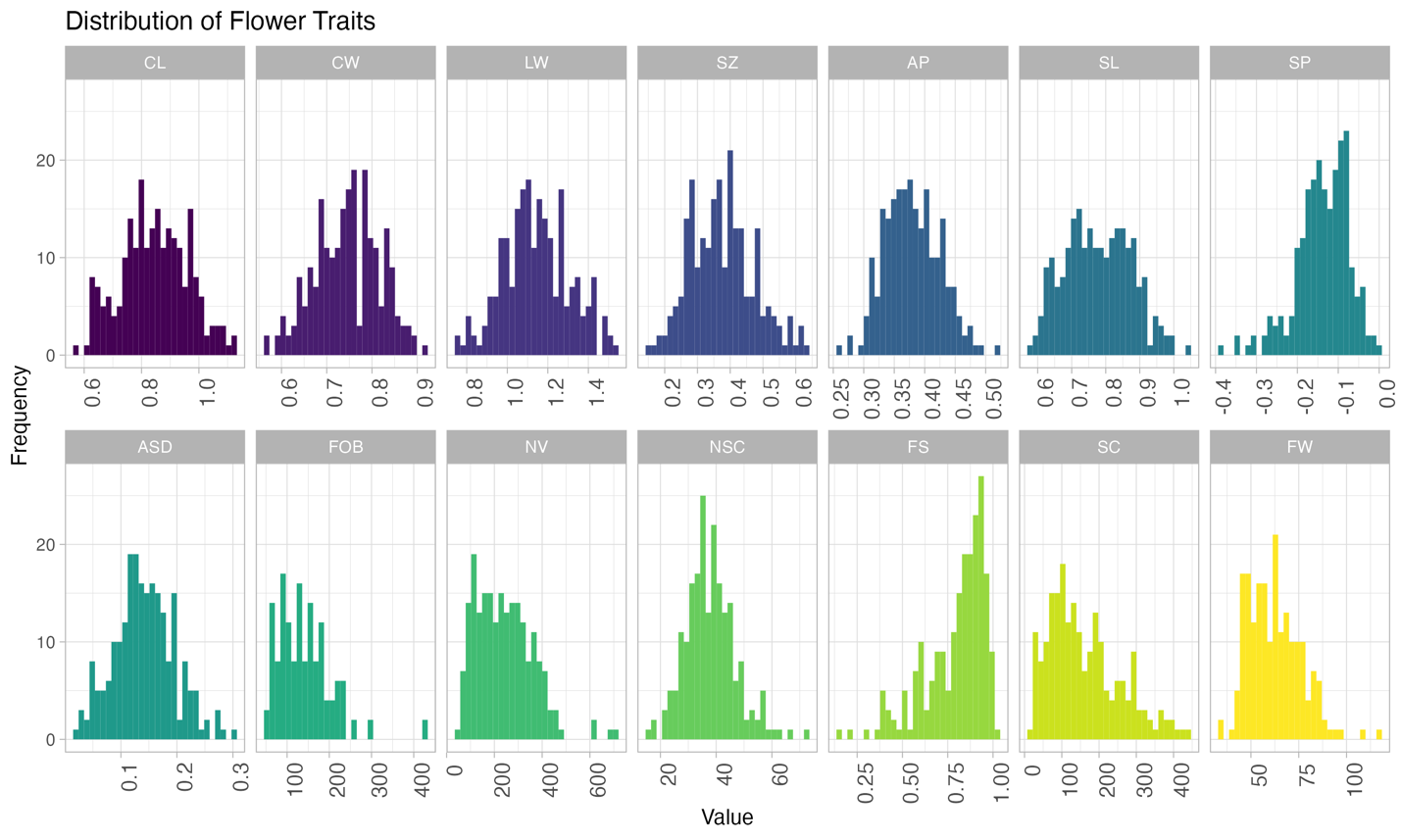

**Supplemental Figure 1.** Histogram distribution of flower morphological traits. CL, corolla length (cm); CW, corolla width (cm); LW, ratio of corolla length-to-width; SZ, flower size (cm^3); AP, aperture diameter (cm); SL, style length (cm); SP, stigma protrusion from corolla (cm); ASD, anther-to-stigma distance (cm); FOB, flowers on bush (flowering density); NV, nectar volume (µL); NSC, nectar sugar content (°BRIX); FS, fruit set; SC, seed count of 10 berries; FW, fruit weight of 25 berries (g).

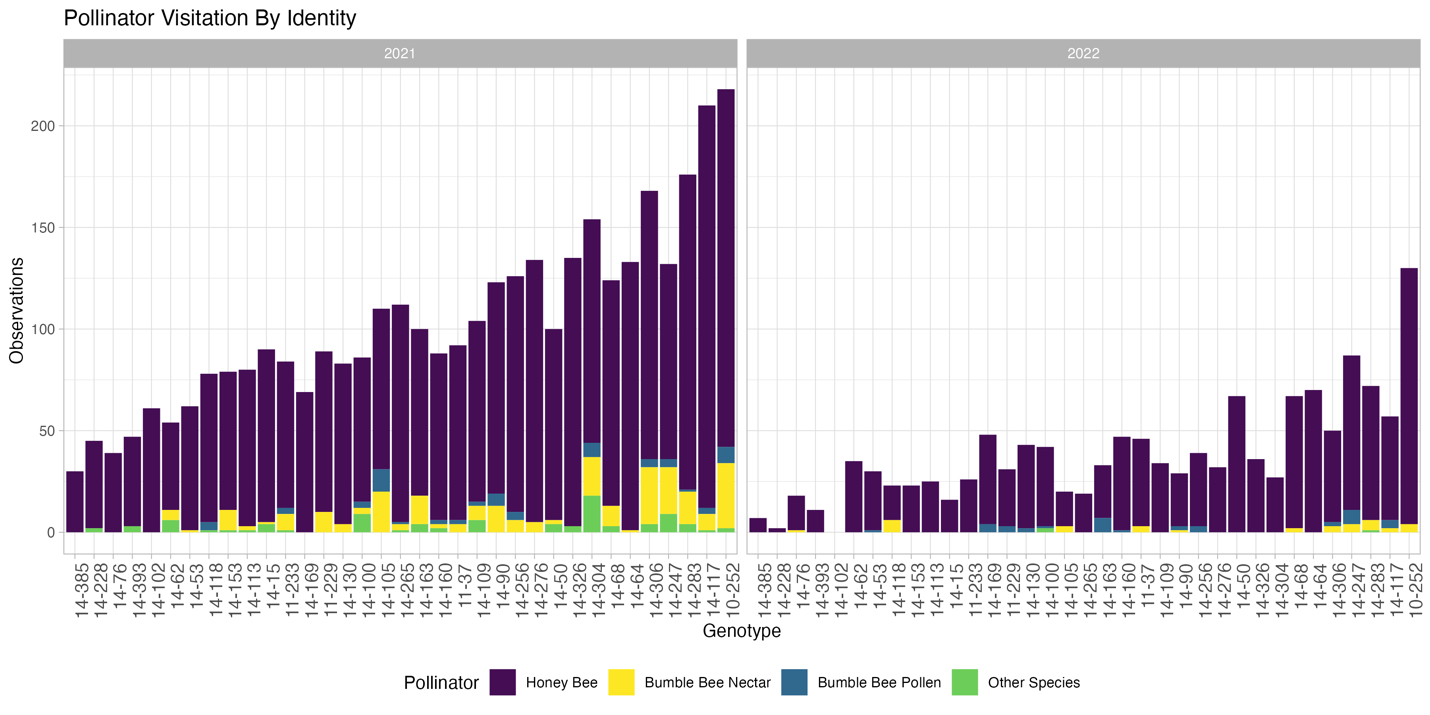

**Supplemental Figure 2.** Pollinator visitation for each genotype between years. Colors indicate pollinator species and foraging behavior. Other species included carpenter bees (*Xylocopa virginica* and *X. micans*), flower wasps (Scoliid spp.), hover flies (Syphridae), and the southeastern blueberry bee (*H. laboriosa*).

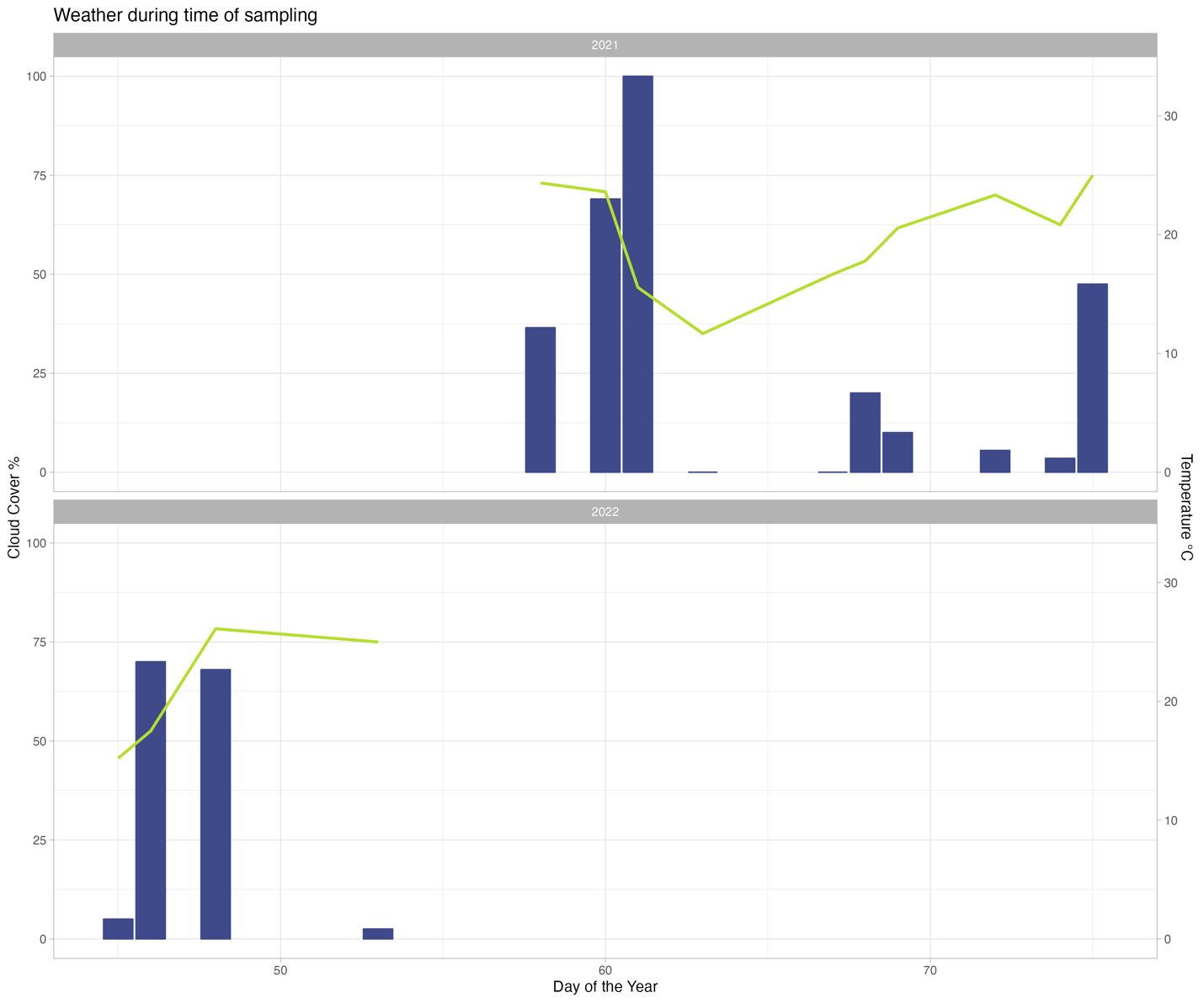

**Supplemental Figure 3.** The temperature (°C) (green line) and cloud-cover percent during the 2021 and 2022 growing seasons. Julian date is presented as day of the year.

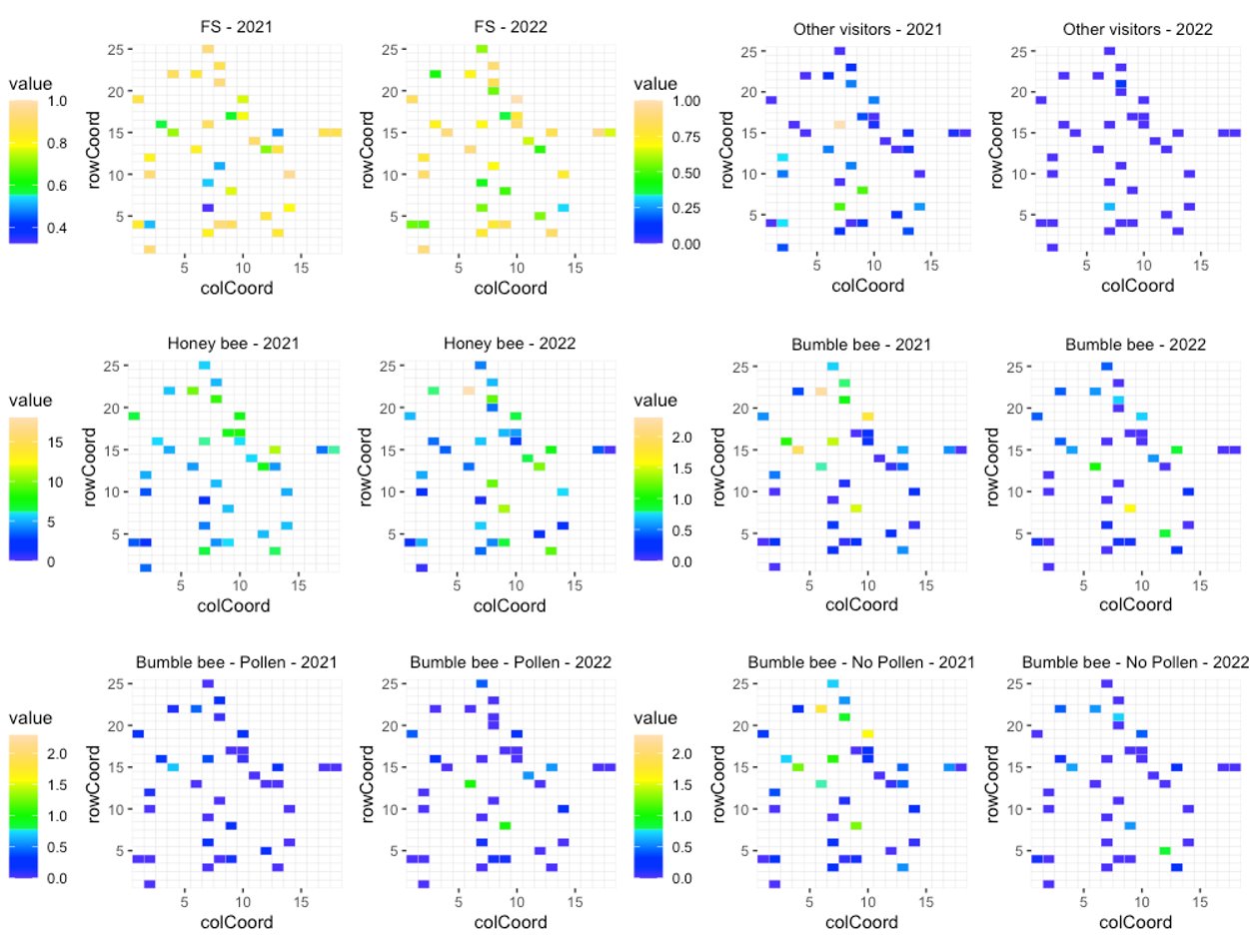

**Supplemental Figure 4.** The spatial distribution of fruit set (FS), honeybee visitation, bumblebee visitation, and other flower visitors across row and column positions in the 2021 and 2022 seasons.

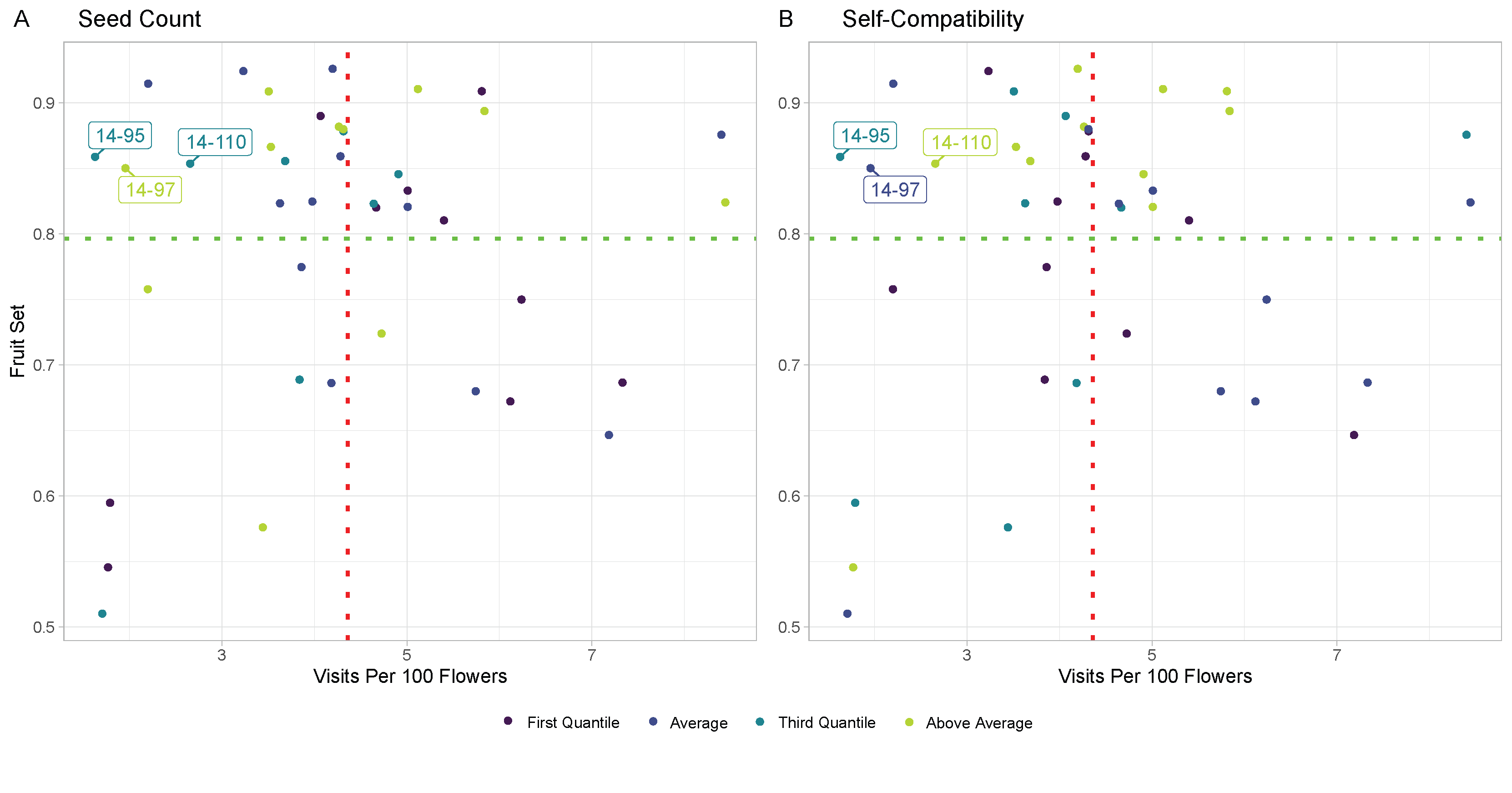
**Supplemental Figure 5.** Outliers removed due to high number of seeds with low pollinator observations (A) and high rates of self-compatibility (B), inflating fruit set at low visitation rates.

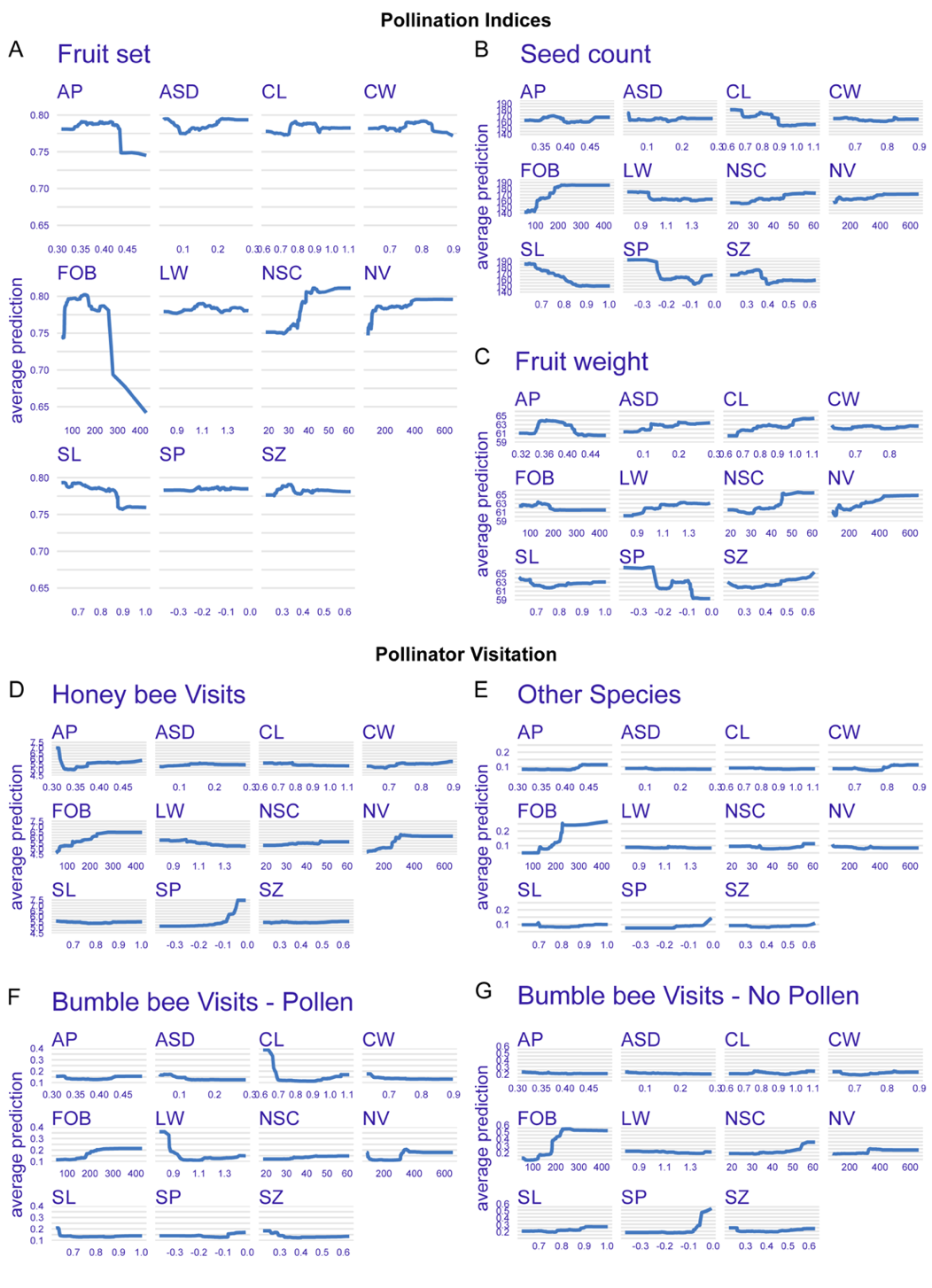

**Supplemental Figure 6.** The partial dependence profile for the relationship between pollination indices (A-C) and pollinator visitation frequency (D-G) with each individual flower trait based on BayesB and Random Forest regression, respectively. AP, aperture diameter; ASD, anther-to-stigma distance; CL, corolla length; CW, corolla width; FOB, flowers on bush (flowering density); LW, ratio of corolla length-to-width; NSC, nectar sugar content; NV, nectar volume; SL, style length; SP, stigma protrusion from corolla; SZ, flower size.
